# Bridging the Data Gap: A Standardized Framework for Monitoring Coral Reefs in Remote Locations via the Recreational Cruising Fleet

**DOI:** 10.64898/2025.12.02.691985

**Authors:** Tom Puchner

**Affiliations:** Planet Ocean

## Abstract

Remote coral reef ecosystems face increasing threats from climate change and anthropogenic pressures, yet comprehensive monitoring remains logistically and financially challenging in isolated regions. This paper presents a standardized framework to bridge this data gap by mobilizing the global recreational cruising fleet as citizen scientists. We describe a low-cost, accessible methodology utilizing widely available action cameras and GPS units to capture continuous video transects without requiring specialized taxonomic training. The collected data is aggregated onto a freely accessible, open-source web platform that integrates high-resolution video with precise geospatial tracking. This system enables remote analysis by experts and machine learning algorithms, facilitating the monitoring of benthic cover, reef health, and bleaching events in areas previously inaccessible to regular scientific survey. By validating a decentralized model for data collection, this framework offers a scalable solution for global reef monitoring while fostering environmental stewardship within the maritime community.

## Introduction

Coral reef ecosystems are currently facing unprecedented peril. Driven by anthropogenic pressures, including climate change, ocean acidification, and overfishing, the rapid degradation of this biome threatens global marine biodiversity and the livelihoods of coastal communities (Hughes et al., 2017; Souter et al., 2021). To effectively manage conservation efforts and understand the trajectory of reef health, there is an urgent need for high-resolution monitoring data that is consistent over time and comprehensive across all tropical latitudes. However, the current scope of data collection is heavily biased toward accessible locations, leaving vast tracts of remote coral ecosystems unmonitored and poorly understood (Roelfsema et al., 2021).

The primary barrier to global monitoring is the logistical and financial burden associated with traditional research methodologies. Scientific expeditions to remote atolls and isolated archipelagos are prohibitively expensive and inherently limited in both temporal frequency and spatial coverage. Furthermore, while community-based monitoring is a valuable tool, remote island communities often lack the necessary technological infrastructure—such as modern high-resolution camera equipment, precise GPS tools, and reliable broadband internet—required to capture and transmit data that meets rigorous scientific standards. Consequently, a significant “data blind spot” exists in the very regions where reefs may be most pristine or, conversely, most vulnerable to unrecorded distinct thermal stress events.

A potential solution to this logistical impasse lies in an untapped global resource: the recreational cruising sailboat fleet. Hundreds of long-distance cruising vessels visit remote tropical locations annually, often anchoring in the precise areas where scientific data is most scarce. Mobilizing these wind-powered vessels not only solves logistical constraints but also aligns with the growing imperative to minimize the carbon footprint associated with marine research and citizen science tourism (Smith et al., 2024). Unlike remote communities that may struggle with equipment costs, most modern cruising vessels are already outfitted with the necessary hardware, including action cameras, reliable GPS navigation systems, and increasingly, satellite-based broadband connectivity. This demographic possesses the mobility and the means to act as effective citizen scientists (De Vargas et al., 2022), yet a cohesive framework to utilize this potential is currently lacking.

To leverage this resource effectively, the monitoring methodology must be robust yet accessible. Video transects represent the optimal approach for this context. Unlike complex point-intercept transects or quadrat surveys, which often require *in situ* taxonomic expertise, video transects allow for the rapid acquisition of data without requiring specialized knowledge of reef ecology or taxonomy. This method not only mitigates observer bias but also creates a permanent visual record that can be analyzed by experts remotely and re-analyzed in the future as diagnostic tools improve. Studies suggest that video transects often yield greater accuracy and archival value than traditional manual survey methods, particularly when conducted by non-specialists (Jokiel et al., 2015).

This paper proposes a novel, standardized framework to mobilize the cruising community for coral observation. We outline a protocol for capturing standardized video transects linked to precise GPS metadata and introduce a freely accessible web-based platform for data submission and visualization. By lowering the barrier to participate and utilizing existing vessels of opportunity, we aim to establish a continuous, decentralized global monitoring network that can provide the essential baseline data needed to protect these critical ecosystems.

## Materials and Methods

### Participant Profile and Training

The data collection methodology is designed to be executed by recreational cruisers with basic to intermediate water skills. No formal scientific training or taxonomic knowledge is required. Participants must be competent snorkelers or freedivers capable of maintaining a steady swimming pace and depth. The protocol relies on “vessels of opportunity”, utilizing cruisers already present in remote anchorages to select survey sites based on local conditions and safety. Survey locations can be suggested remotely based on previous surveys or publicly available aerial imagery.

### Equipment and Configuration

To ensure data quality and standardization across different observers, the following equipment specifications are required:

- **Imaging Device:** A waterproof action camera (e.g., GoPro or equivalent) capable of 4K recording. Image stabilization features are highly recommended to facilitate accurate post-hoc analysis.
- **Camera Settings:** Video resolution is set to 4K at 30 frames per second (fps). The “Linear” zoom setting is mandatory to minimize fisheye distortion, which can compromise areal measurements. If hardware constraints exist, reducing the frame rate is preferred over reducing resolution (minimum acceptable resolution is 2.7K).
- **Geolocation:** A continuous track recording is required via a mobile GPS unit, GPS-enabled watch, or a smartphone running a tracking application housed in a waterproof bag/case.
- **Accessories:** A swim buoy with a tether or a waterproof bag to secure the GPS unit to the swimmer.

#### Synchronization Protocol

Prior to deployment, the internal clocks of the action camera and the GPS device must be synchronized to the nearest second. This step is critical for the subsequent temporal correlation of video frames with geographic coordinates.

### Field Protocol: The Video Transect

The survey utilizes a timed swim method to approximate a 20-meter transect. The GPS tracking is initiated prior to water entry to ensure a continuous satellite lock. Multiple transect videos can be recorded during one in-water-session (Fig. 1). The continuous GPS track is later divided and assigned to the single video files. Start and endpoint of each transect GPS track is determined by either of three methods: Synchronized time, 90° angle and several track points recorded at the same location.

**Figure 1.**
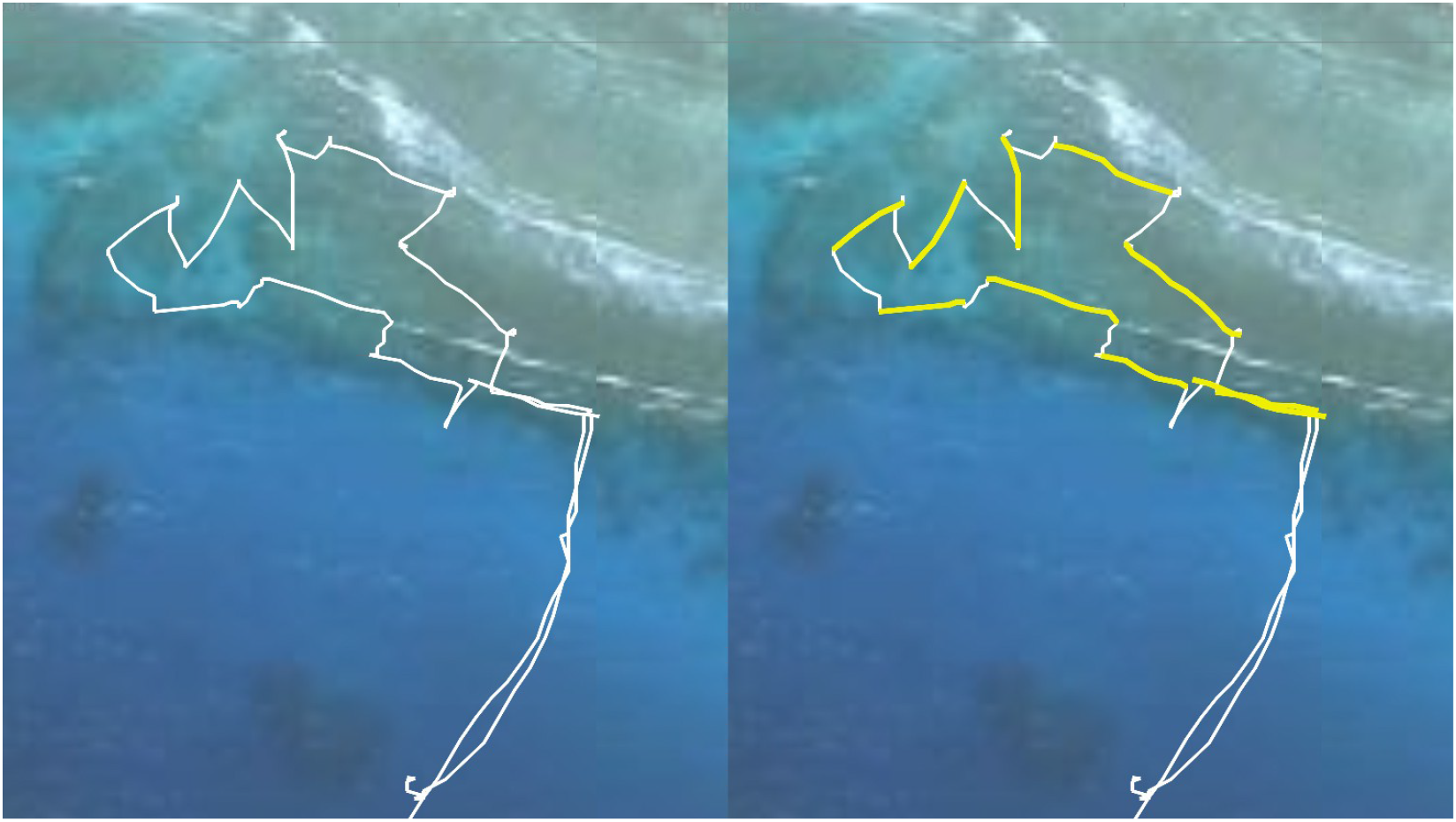
Segmentation of GPX track into individual transects. The left panel displays the raw, continuous GPX track of an entire in-water session. The right panel illustrates the segmentation process, where individual transects (highlighted in yellow) are delineated from the continuous track by using the distinct 90° turns made by the observer during the Approach and Egress phases as start and end markers. These transect tracks with associated videos can be viewed at https://www.planet-ocean.org/corals.html?center=177.052330,-17.590729&zoom=19&layer=arcgis

**Figure 2.**
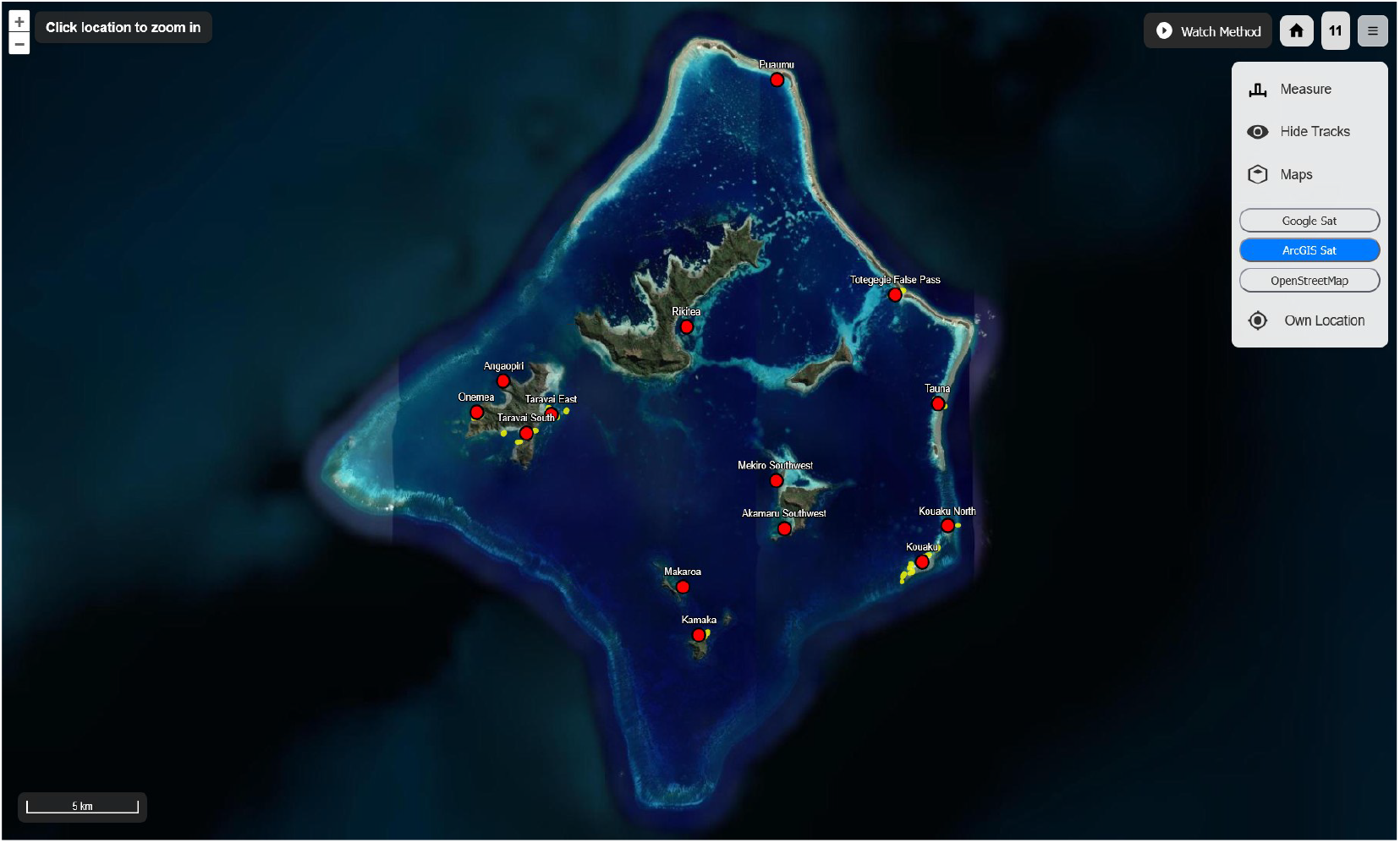
Web Application Visualization at Region Level (Zoom Level 9-11). The satellite map displays the Gambier Islands in French Polynesia overlayed with monitoring data. Red dots represent transect locations; yellow lines indicate individual transects. The user interface elements for layer selection and measurement are visible on the right.

**Figure 3.**
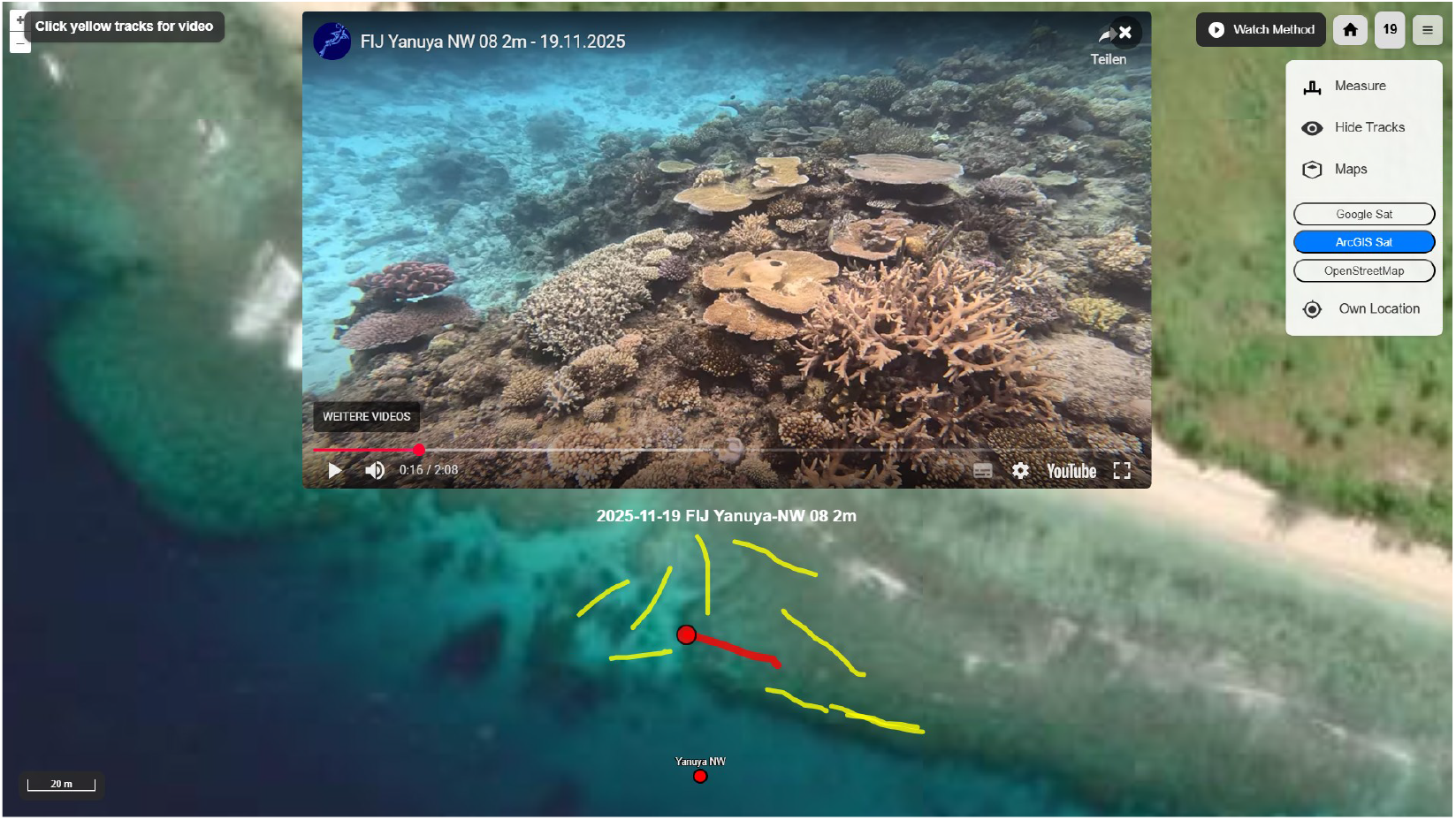
Web Application Visualization at Location Level (Zoom Level 16-19) with Video Playback. This high-resolution view shows the individual track data overlayed on the satellite imagery. The red line represents the GPS track of the active transect with video popup, while the yellow lines show all available transects at the location. Clicking a yellow track triggers a popup (top of image) that instantly embeds and plays the associated survey video, providing immediate visual access to the collected benthic data. These transect tracks with associated videos can be viewed at https://www.planet-ocean.org/corals.html?center=177.052285,-17.590657&zoom=18&layer=arcgis

#### Transect Execution

- **Approach:** The observer identifies a safe starting point and approaches the transect start location at a 90° angle (perpendicular to the intended swim path). This geometric approach aids in pinpointing the exact start location on the GPS track during analysis.
- **Recording Sequence:**
  - **00:00 - Surface Orientation:** The recording begins above water, facing the direction of the swim to capture coastal landmarks or horizon context.
  - **00:02 - Subsurface Context:** The camera is submerged to slowly capture a 360° panoramic view of the surrounding reef environment.
  - **00:30 - Metadata Signaling:** The observer indicates depth using hand signals in the field of view (e.g., three fingers for three meters). For depths under one meter (reef flats), a bent index finger is used. If the topography is sloping, maximum and minimum estimated depths are indicated.
  - **Transect Swim (00:35 – 01:35):** If the depth exceeds 2 meters, the observer dives to about 1.5m above the bottom. The observer swims slowly in a straight line for approximately 60 seconds (covering ~20 meters), maintaining a camera-to-bottom distance of approximately 1 meter. The camera is held facing downwards and slightly forward. Deviations from a straight line are permitted to follow depth contours.
  - **Termination (01:35 – 02:05):** The observer ascends to the surface, tilts the camera horizontally for a final slow 360° subsurface view, and stops recording.
- **Egress:** The observer swims away from the end point at a 90° angle to clearly demarcate the transect finish on the GPS track.

#### Adaptations for Shallow Environments

For reef flats accessible at high tide, diving is not required. The observer remains at the surface, tilting the camera forward to maintain the requisite 1-meter distance from the benthos while ensuring the sensor remains submerged.

### Data Management and Submission

Following the survey, data integrity is maintained through a standardized file hierarchy. Participants create a specific directory named according to the format: <monospace>YYYY-MM-DD - YYYY-MM-DD [Location Name], [Vessel Name]</monospace>. Both the raw video files (.MP4/.MOV) and the complete GPS track (preferably in.GPX format) are deposited into this directory. In the pilot phase, data submission is conducted via cloud storage links, direct email, or upload to designated video hosting platforms. Future iterations of the project will utilize a dedicated web interface featuring automated upload and synchronization capabilities.

## Results

At the time of writing, a total of 343 transects have been conducted, mainly in French Polynesia and Fiji. To estimate the benthic surface area surveyed, we calculated the camera’s effective field of view (FOV) underwater. The GoPro “Linear” digital lens setting provides a horizontal FOV of approximately 86° in air; however, the activation of “HyperSmooth” stabilization introduces a crop factor of approximately 10%, reducing the effective air FOV. Furthermore, applying Snell’s law of refraction (n1sinθ1=n2sinθ2) with the refractive indices of air (n≈1.0) and seawater (n≈1.34) through a flat port housing significantly narrows this angle. The resulting effective underwater horizontal FOV is approximately 56°. With a standardized camera-to-bottom distance of 1 meter, this geometry yields a transect width of roughly 1.1 meters. Consequently, each 20-meter transect covers approximately 22 m^2^, bringing the approximate total area covered by the 343 transects to 7,500 m^2^.

### Web Application and Visualization Platform

The immediate operational outcome of this framework is a centralized, freely accessible web application, available at **www.planet-ocean.org/corals.html**. This platform aggregates submitted data and provides an interactive interface for global visualization of the monitoring network.

#### Technical Architecture

The application is built upon the OpenLayers library, ensuring robust performance and compatibility across devices. To facilitate precise geographic context, the map interface supports multiple base layer overlays, including ArcGIS, Google Satellite, and OpenStreetMap. This flexibility allows users to cross-reference transect data with satellite imagery for habitat verification.

#### Data Representation

Survey data is visualized spatially. The GPX track of each individual transect is rendered as a vector line directly on the map. The user interface is designed for intuitive interaction; clicking on a specific transect line triggers a popup window containing the associated video footage, allowing for immediate visual assessment of the benthic cover at those specific coordinates.

#### Navigation and Scalability

To manage the disparity between global coverage and local detail, the interface employs a hierarchical navigation system. The default view presents a global perspective (Zoom Level 2). Users can seamlessly navigate through three distinct scales:

- **Large Marine Ecosystem/Country Level (Zoom Level 6):** For broad regional assessment.
- **Region Level (Zoom Level 9-11):** For archipelago or island-group specific analysis.
- **Location Level (Zoom Level 16-19):** For high-resolution viewing of individual transects and reef topography.

#### Analytical Tools and Field Utility

The platform includes integrated tools to support both remote analysis and in-field operations. A measuring tool allows researchers to verify transect lengths or measure other spatial features directly on the map canvas. Furthermore, the application supports geolocation (displaying the user’s current position), a critical feature for cruisers attempting to revisit specific transects for longitudinal monitoring. Finally, the system supports deep linking; clicking on the web canvas generates a unique URL for the current map view, facilitating easy sharing of specific reef locations and datasets among researchers and the citizen science community.

## Discussion

### Analysis Architectures: From Volunteers to Algorithms

The high-resolution video data collected through this framework supports a flexible, multi-tiered analysis pipeline adaptable to varying resource levels and scientific requirements. At the most basic level, the raw footage serves as a qualitative archival record. However, for quantitative monitoring, the standardized nature of the transects facilitates processing by diverse analytical actors. Trained volunteers and citizen scientists can annotate footage for coarse functional groups, while taxonomic experts can utilize the same dataset for genus- or species-group-level identification. Crucially, the standardized 4K imagery is ideally suited for processing by emerging machine learning (ML) algorithms (Sauder et al., 2023; Ouassine et al., 2025). By feeding this geographically diverse dataset into ML models, we can accelerate the extraction of benthic cover metrics, making near-real-time global monitoring a tangible possibility.

### Implications for General Reef Monitoring

The primary value of this decentralized network lies in its ability to generate consistent time-series data for otherwise data-deficient regions. The video transects allow for the robust quantification of essential ecological metrics, including percentage of hard coral cover, substrate composition (bottom type), and dominant growth forms. Beyond static cover metrics, the high frame rate and resolution enable the assessment of dynamic reef health indicators. This includes the early detection and mapping of mass bleaching events, tracking the prevalence of coral diseases, and monitoring recruitment and recovery following disturbances. Over extended periods, this data is vital for identifying subtle phase shifts—such as the transition from *Acropora*-dominated to *Pocillopora*- or *Porites*-dominated states—and changes in species composition that might otherwise go unnoticed in remote locations (Speelman et al., 2023; Cannon et al., 2021).

### Supporting Coral Restoration Initiatives

The utility of this framework extends beyond passive observation to active restoration support. The combination of the web platform’s aerial imagery integration and the “ground-truth” video data provides a powerful tool for remote site prospecting. Restoration practitioners can identify potential sites for genotype collection, nursery establishment, or outplanting without the need for initial, expensive exploratory expeditions (Bowden-Kerby, 2025). Furthermore, the platform enables remote support for local island communities interested in establishing their own nurseries but lacking technical baseline data. Once restoration projects are established, the cruising fleet can provide ongoing monitoring support, visiting these sites seasonally to document growth rates and survival, thereby ensuring the long-term viability of restoration efforts.

### Immersion and Inspiration

As long-term cruisers with over 13 years of experience and more than 75,000 nautical miles navigated across tropical seas, we are uniquely positioned to initiate this undertaking. This extensive immersion grants us direct access to the global cruising community, facilitating the peer-to-peer engagement necessary to sustain such a decentralized network. The impact of this project transcends the data itself. By integrating the cruising community into the scientific process, we foster a deeper sense of immersion and stewardship among ocean users. The act of conducting a transect transforms a passive visitor into an active participant in reef conservation. This experiential learning has the potential to inspire a broader culture of environmental responsibility within the maritime community, creating a fleet of knowledgeable advocates for the marine ecosystems they traverse.

